# Impact of Zygosity in Bimodal Phenotype Distributions

**DOI:** 10.1101/086215

**Authors:** Thomas Holst-Hansen, Elena Abad, Aura Muntasell, Miguel López-Botet, Mogens H. Jensen, Ala Trusina, Jordi Garcia-Ojalvo

## Abstract

Allele number, or zygosity, is a clear determinant of gene expression in diploid cells. But the relationship between the number of copies of a gene and its expression can be hard to anticipate, especially when the gene in question is embedded in a regulatory circuit that contains feedbacks. Here we study this question making use of the natural genetic variability of human populations, which allows us to compare the expression profiles of a receptor protein in natural killer cells between donors infected with human cytomegalovirus (HCMV) with one or two copies of the allele. Crucially, the distribution of gene expression in many of the donors is bimodal, indicative of the presence of a positive feedback somewhere in the regulatory environment of the gene. Three separate gene-circuit models differing in the location of the positive feedback with respect to the gene can all reproduce well the homozygous data. However, when the resulting fitted models are applied to the hemizygous donors, only one model (the one with the positive feedback located at the level of gene transcription) reproduces the experimentally observed gene-expression profile. In that way, our work shows that zygosity can help us relate structure and function of gene regulatory networks.

**Author Summary:** Nearly all mammalian cells, including human cells, have two copies of each chromosome, and thus possess two potentially different copies of each gene (which might be in some cases non-functional or even absent). Naïively one might expect that two identical copies of the gene would lead to the protein being expressed at twice the rate, but many factors can alter this simple calculation. One of these factors is the existence of feedback mechanisms affecting in one way or another the regulatory circuit in which our gene of interest is embedded. Here we study the relationship between the number of gene copies and the expression of a receptor protein that plays a crucial role in the recognition of pathogens by natural killer cells, which are important elements of the innate immune system. Experimental data of virus-infected donors reveals a bimodal expression profile of this receptor, typical of a positive feedback, and a clear difference between donors with one or two copies of the gene. Mathematical modeling allows us to find the likely location of the feedback loop within the gene’s regulatory circuit, by requiring the correct model to reproduce the expression profiles of both types of donors.

## Introduction

Natural killer (NK) cells are part of the innate immune system, with the primary role of detecting and eliminating virally infected cells and tumours. They recognize potential threats through a gauging process, where signals from activating and inhibitory receptors are integrated into a decision. A target cell is killed either if inhibitory receptors are not engaged, or if the NK cell receives a strong activating stimulus. Inhibitory receptors sense MHC class-I surface molecules that present intracellular peptides [1]. Since MHC molecules can be lost due to mutations in tumors or viral infection, NK cells are particularly important in the defence against those threats [2, 3].

The innate immune system provides a generic response to pathogens, and contrary to the adaptive immune system, it does not create long-term immunity. However, NK cells (which as mentioned above are considered innate immune cells) share adaptive memory features with T-cells in response to certain viruses [4]. For instance, in response to infection with human cytomegalovirus (HCMV), some donors experience stable expansion of the NK subset containing the activating NKG2C receptor [5–7], resembling the memory of the adaptive immune system.

HCMV is a herpes virus that establishes a persistent infection within the host and is extremely common, ranging from 45%-75% prevalence in Western countries to close to 100% in developing countries [8]. The virus has an elaborate arsenal of evasion strategies that protect it from recognition, such as downregulation of MHC-I and expression of MHC-I decoys [9]. The NKG2C+ NK cell subset is believed to play a role in antiviral defense, and has been shown *in vitro* to mediate a potent antibody-dependent response against HCMV-infected cells [10]. The expansion of this NK cell subset involves different elements of the immune system, including IL-12, CD14+ cells (e.g. macrophages and dendritic cells) and the CD94/NKG2C/HLA-E axis [11, 12]. But also zygosity (gene copy number) affects the NKG2C prevalence. Specifically, homozygous (two gene copies) and hemizygous (one gene copy) donors differ significantly in the expression level and function of the NKG2C receptor, as well as in the fraction of NK cells that express this receptor [13].

Notably, HCMV-infected donors usually expand the NKG2C+ NK cell subset in a bimodal fashion [13]. Bimodal distributions are indicative of a bistable response, which is commonly generated by positive feedbacks [14]. In this paper we show that three types of positive feedback, located at different positions with respect to the NKG2C gene (upstream of the gene transcription, directly at the level of gene transcription, and at the post-transcriptional level) can generate similar bimodal distributions in homozygous donors. However, a comparison with the corresponding hemizygous distributions allows us to differentiate between these three cases. Specifically, our computational model shows that the distributions of NKG2C+ NK cells are best described by a positive feedback at the level of NKG2C transcription, as opposed to positive feedbacks located pre- or post-transcriptionally. We use bifurcation analysis to show mathematically that zygosity leads to two types of changes in the bistable expression distributions: changes in expression level, and changes in the fraction of cells expressing the phenotype. As we will see, a positive feedback upstream of the gene transcription leads only to expression level changes, while a post-transcriptional positive feedback just leads to changes in the fraction of expressing cells. In contrast, a positive feedback acting directly at the level of gene transcription has both effects in response to zygosity changes: in the expression level and in the fraction of expressing cells.

Copy number variations (CNV) are a common form of genetic variability, and are linked to various autoimmune diseases such as rheumatoid arthritis [15], asthma [16], and susceptibility towards HIV [17]. However, the link between CNVs and phenotype expression can be elusive. The mathematical framework presented here can serve as a tool for understanding the interactions between gene copy number and the topological architecture of cellular regulatory networks.

## Results

### Effect of zygosity on NKG2C expression distributions: experimental observations

We quantified NKG2C levels using flow cytometry. The profiles of many of our HCMV-positive donors are clearly bimodal (Fig 1A,B). Data from nullzygous donors (homozygous for the NKG2C deletion, Fig 1C) shows that the left peak of the bimodal distribution is purely background fluorescence, since these individuals do not express the receptor. Based on the nullzygous donor profiles we define an expression threshold of *R* = log_10_(NKG2C) = 2.75 (leftmost vertical dashed line in Fig 1A-C). Cells are termed ‘expressing’ if they have an expression level above that threshold (Fig 1D). The fraction of cells that express NKG2C is calculated as the area under the normalized NKG2C distribution beyond the expression threshold. Similarly, the average expression level is only calculated for cells above the expression threshold.

The data reveals a significant decrease in both the expression level (Fig 1E) and fraction (Fig 1F) of NKG2C+ NK cells, when comparing homozygous with hemizygous donors. In the two cases the distribution of NKG2C in the NK population of individual donors is bimodal, but also highly variable between donors.

**Fig 1.**
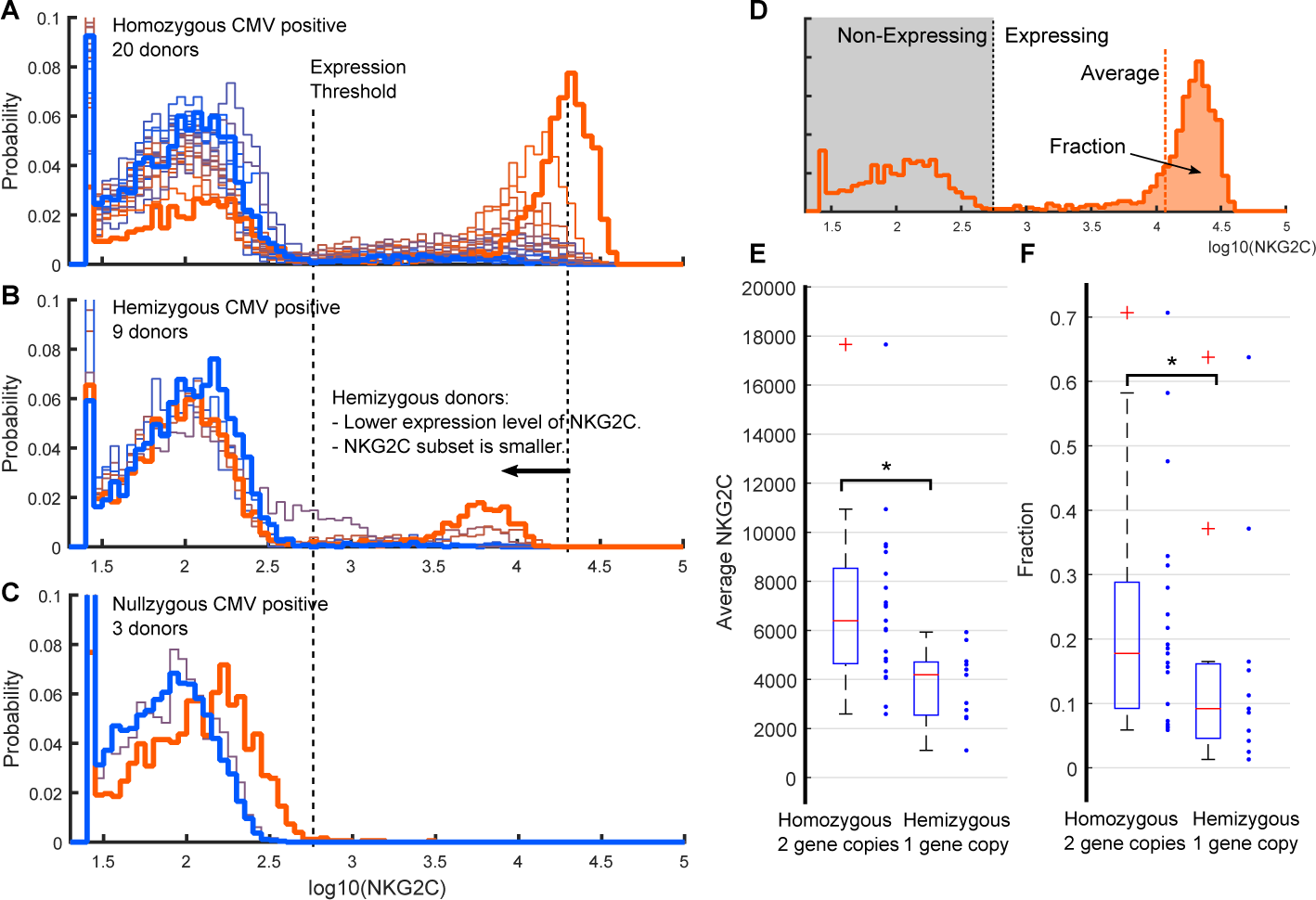
Zygosity influences expression level and prevalence of NKG2C in NK cells of HCMV positive donors. Flow cytometry measurements of NKG2C in NK cells [13], for HCMV positive homozygous (**A**), hemizygous (**B**), and nullzygous (**C**) donors. Individual donors are color-coded according to the NK population fraction expressing the NKG2C receptor, with orange (blue) corresponding to the highest (lowest) expressed fraction. Panel **D** shows how the distributions are quantified. Specifically, the fraction of expressing cells is defined as the area under the normalized distribution above the expression threshold. Likewise, the average expression level of cells is calculated for those cells above the expression threshold. Panels **A** and **B** show that reducing gene copy number lowers both the expression level and the fraction of NK cells expressing NKG2C. Panels **E, F** show box plots quantifying the significant differences between homozyogus and hemizygous donors in the average NKG2C level of expressing NK cells, and in the fraction of the NK population expressing NKG2C. In those two panels, the asterisk indicates a statistically significant difference, using a two-sided Mann-Whitney U test with significance level of *p* = 0.05 (see Supporting Table ??).

The first key assumption in the comparison of the model with the data is that the fluorescence measurements are proportional to the surface level of NKG2C [18]. Second, the measurements should be representative of the total NK population and be in a state of homeostasis. Third, the steady-state levels of NKG2C+ NK cells remains stable in healthy donors [13, 19]. Further, it should be specified that measurements are a snapshot of the peripheral NK population, as they are made on blood samples.

### Modeling NKG2C bimodality

The aim of the model is to describe the impact of zygosity (gene copy number) on the expression level of the NKG2C receptor in NK cells, and on the fraction of cells in the population expressing this receptor. A key observation that constrains the model is that the population is bimodal in its expression of NKG2C. We assume that this bimodality is caused by a positive feedback. We consider three models that differ on where the positive feedback is located with respect of NKG2C expression (Fig 2A). In model **A** the feedback occurs upstream of NKG2C expression, at the level of a transcription factor regulating the expression of the gene (named pre-transcriptional in what follows). In model **B** it arises at the level of transcription of the gene. Finally, in model **C** the feedback is considered to occur at the post-transcriptional level. All three models describe the dynamics of three variables: a transcription factor *T*, the mRNA *m*, and the mature receptor *R*. The model is kept minimal, in order to illustrate the effects of bistability and gene copy number, and as such it does not take into account specific proteins or processes. The variables could therefore correspond to any part of the signalling pathway with a positive feedback onto itself.

**Fig 2.**
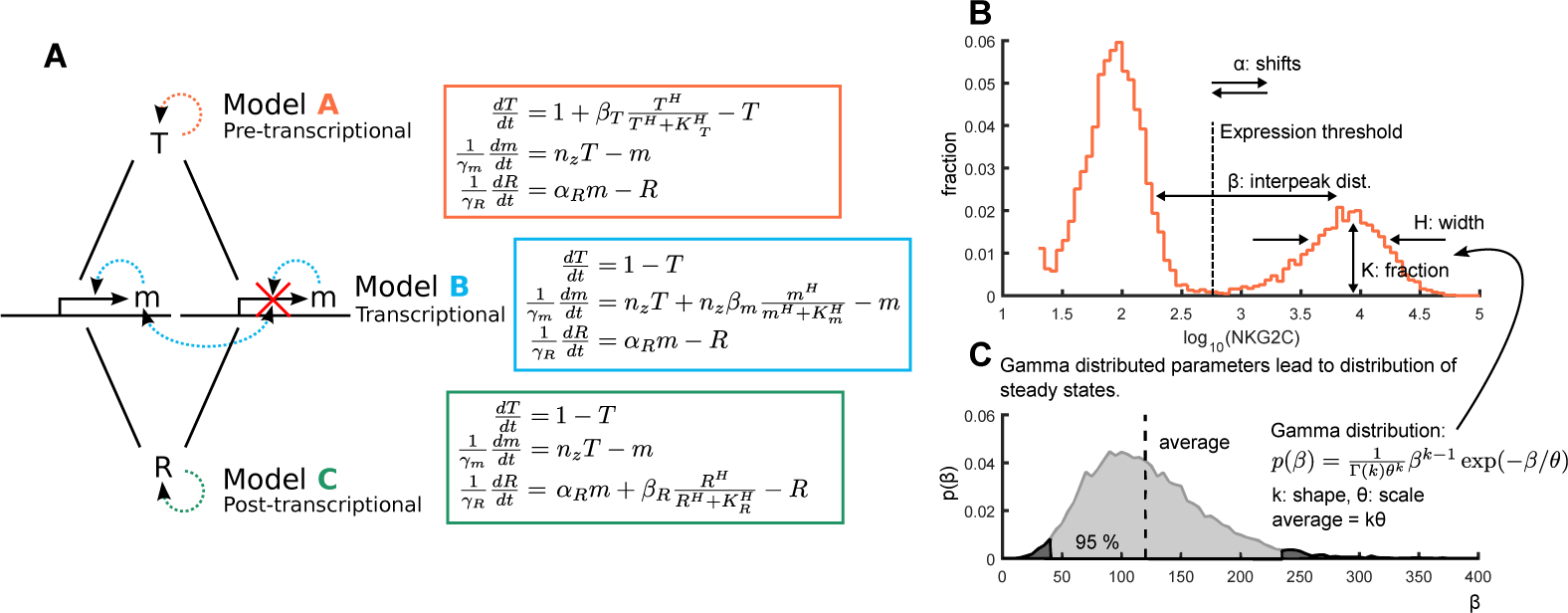
Three minimal models of NKG2C expression. **A**: The three models differ in the position of the feedback relative to transcription; A, a pre-transcriptional, B, a transcriptional and C, a post-transcriptional positive feedback. Each model is described by three differential equations governing the dynamics of *T*, a transcription factor, *m*, mRNA concentration, and *R*, mature receptor. The parameter *n*_*z*_ denotes the gene copy number in each model, and the positive feedback is incorporated as a Hill term. **B**: Each model parameter affects the shape of the overall distribution in different ways: *α* shifts the entire distribution (only model A and B), *β* changes the distance between the two peaks, *K* controls the fraction of cells in each peak and *H* the sharpness of transition between peaks, i.e. the width of the peak and especially the region between the two peaks. **C**: The distribution of the NKG2C receptor (*R*) is generated by drawing the parameters *α, β, K* and *H* from gamma distributions.

In model **A**, for instance, *T* has a positive feedback onto itself implemented with a Hill term. For simplicity we describe production and degradation as linear terms:

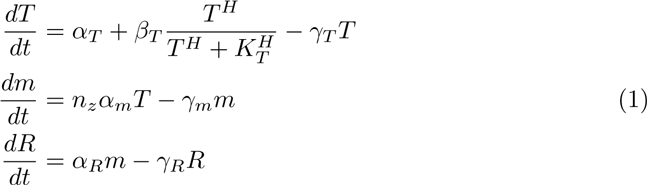

Here *α*_*T*_ is a basal production, *α*_*m*_ and *α*_*R*_ are linear production rates, *β*_*T*_ is the strength of the positive feedback, *H* is the Hill coefficient, *K*_*T*_ is the activation threshold and *γ*_*x*_ are degradation rates. *n*_*z*_ is an integer which corresponds to gene copy number. Note thatthemodeldescribesthereceptorlevel,whilethedonordistributionsaremeasures of fluorescence from flow cytometry. This means that the first peak in the experimental measurements actually corresponds to background activity in the absence of receptors. In the model we reproduce this activity through the basal production coefficient *α*_*T*_. Each model has nine parameters, but by rescaling we reduce the number of variables to six (see Supporting Information). Models **B** and **C** are similar to model **A** above, with the positive feedback on different variables. The rescaled equations are shown in the right panels of Fig 2A. We have not scaled *R* in terms of *α*_*R*_, as we want to maintain that parameter as a fitting parameter.

We assume that the NK population of donors is in homeostasis at the time of measurement. Therefore we only consider steady states, which for the rescaled version of model **A** are given by the following equations:

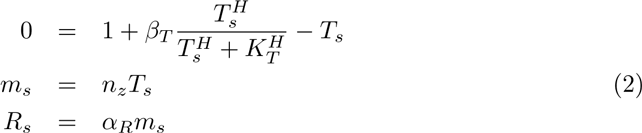

For *T ≫ K*_*T*_, the first equation reduces to *T*_*s*_ = 1 + *β*_*T*_, while in the regime where 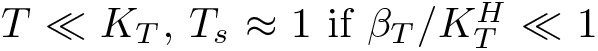, which is a good approximation in our case because the majority of cells are not expressing the receptor. This approximate solution shows that the separation between peaks is controlled by the *β* parameter (Fig 2B). The Hill coefficient *H*, in turn, controls the width of the peaks, since a higher coefficient leads to a sharper transition between solutions. Finally, the threshold *K*_*T*_ of the positive feedback regulates the fraction of cells expressing the receptor, and *α* scales both solutions simultaneously. The same is true for model **B**, while in model **C**, *α* only controls the position of the first peak. Note that the steady states also predict no receptor production in the case of nullzygous donors.

Noise is needed to generate a population of diverse cells. We assume that differentiated NKG2C+ NK cells do not switch receptor expression during their life-span, so that the bimodality occurs only at the population level. Therefore we have chosen to introduce noise by drawing parameters from distributions, rather than adding dynamical noise via stochastic simulations, as is commonly done when studying stochastic gene expression. The abstract character of our model also makes stochastic simulations less suitable, since intermediate steps that could introduce noise are not made explicit. Each parameter is drawn from a gamma distribution (Fig 2C), described by a shape coefficient *k* and a scale coefficient *θ*, so that the average of the distribution is *kθ*. The gamma distribution is a good description of the statistics of protein abundance [20]. In the Supporting Information we show how noise in each parameter affects the distribution. The distribution of NKG2C expression in the NK population is simulated by drawing a set of parameters for each cell. Each of the four parameters defining the steady state (Eq 2) is drawn from a distribution with given average and scale coefficient, which gives eight free parameters in total. The NKG2C distributions were generated by solving numerically the nonlinear equations of each model, and choosing the lowest solution in those cases where there were two stable solutions (since the bistable region is narrow compared with the distribution of parameters, the difference between choosing the low and high solution in the bistable region is small).

### Transcriptional Positive Feedback Best Captures Zygosity Impact in HCMV Positive Donors

To compare each of our models with the experimentally measured NKG2C expression levels of the homo- and hemizygous HCMV positive donors (Fig 1), we calculated a profile of the “median donor” for each group, as the median frequency of the cell number at each expression level. We then fitted each model to the homozygous donor group and achieved fits of similar quality (measured in terms of the *χ*^2^-value), as shown in Fig 3A (see Methods section and Supporting Information for fitting procedure). Hence, it is not possible to differentiate between the three models based only on homozygous donors.

**Fig 3.**
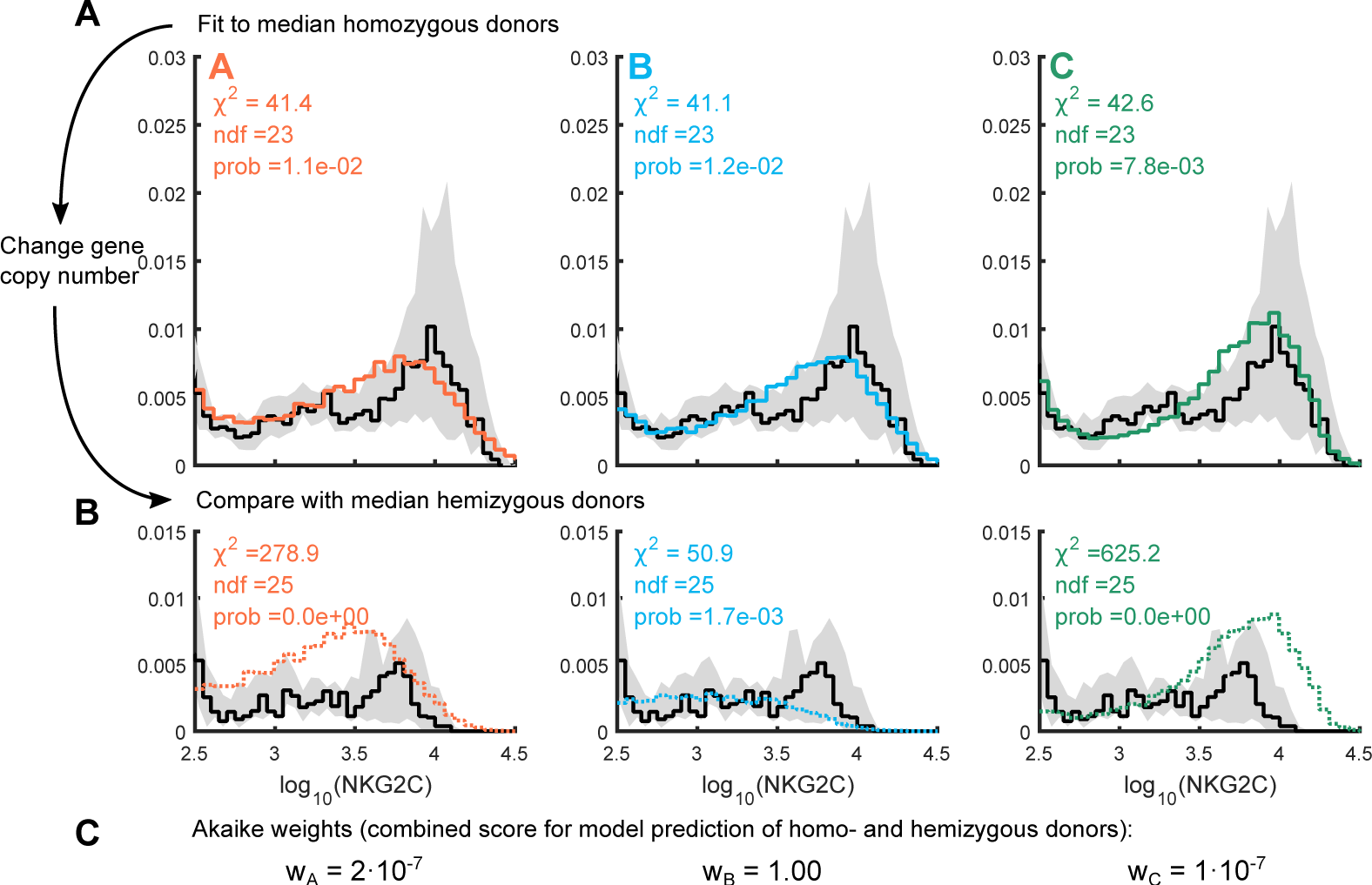
Model B makes the best prediction of NKG2C expression in hemizygous donors. **A**: Each model is fitted to a median homozygous donor (black lines), generated by taking the median cell frequency at each receptor level. The grey area shows the 50% interquartile region. The model distributions provide fits of similar quality (color lines) based on a Pearson’s *χ*2-value. **B**: Starting from the fit of the median homozygous donor, the gene copy number is changed from two to one, and the similarity between the median hemizygous donor and the models are scored with a *χ*2-value. **C**: The Akaike weights are calculated based on both the homo- and hemizygous simulation, and are indicative of the quality of each model in fitting the data, compared with the other models. Model B is clearly favoured over both A and C. All distributions were generated from 100.000 simulated cells.

Using the parameters found by fitting the median homozygous donor, we next changed the gene copy number *n*_*z*_ from two to one. Each model makes different predictions of the median NKG2C distribution of hemizygous donors, which makes it possible to distinguish between models (Fig 3B). In particular, model **B** provides the best prediction of the hemizygous donor group, indicating that a positive feedback at the transcriptional level is the most likely of the three possibilities.

The models we present here are simple and do not capture some details of the distributions. Thus, as an measure of relative model quality alternative to the *χ*^2^-value, we also used the Akaike information criterion (AIC). The Akaike weights, shown in Fig 3C, are based on the combined model prediction of homo- and hemizygous donors. Again, model **B** is clearly the most likely of the three models in describing the NKG2C changes due to zygosity. Biologically, this means that an upstream or downstream positive feedback is not sufficient to describe, at the single cell level, the impact of zygosity on NKG2C expression.

From a modeling point of view the virus has the effect of increasing the strength of the positive feedback, *β*, causing higher levels of NKG2C in the NK population. Secondly it lowers the activation threshold, *K*, of the positive feedback, which increases the fractional expression of NKG2C. This could occur for instance through promoter alterations or increased receptor activation. But in any case, our results show that any positive feedback should start and end at the transcriptional level, as described by model **B**.

### Bifurcation Diagrams Present Model Differences

We can use bifurcation diagrams to translate a distribution of parameters into a distribution of NKG2C expression. Changing the model zygosity will alter the bifurcation diagram, and therefore the corresponding receptor expression. In Fig 4 we show the bifurcation diagram of each model for varying activation thresholds *K*, and depict how a distribution of *K* is translated into receptor expression. Our analysis shows that all three models undergo two saddle-node bifurcations, delimiting the region in which three stable fixed points exist (two of them stable, represented by solid lines, and one unstable, depicted by a dashed line). For low activation thresholds a single stable solution exists at high NKG2C levels. As the activation threshold increases, the low stable solution and an unstable solution is created. Further increase of *K* causes the unstable and the high stable solution to annihilate. The position of the bifurcation determines the fraction of the *K*-distribution that is translated into expressing cells, while the value of the solution corresponds to the NKG2C expression level. We adjust the position and width of the parameter distributions relative to the bifurcations so as to recover as closely as possible the experimentally observed receptor distributions.

In model **A** the bifurcations are positioned at the same *K*-values independent of zygosity, while the solutions are shifted by approximately a factor two. This results in a change in expression level, but not in the fraction of cells expressing the receptor. In contrast, in model **C** zygosity causes a shift in the first bifurcation while the other remains the same. This lead to a changed fraction of expressing cells that is too small to reproduce the experimental observations. The effect of zygosity on the high stable solution is negligible, which causes the position of the second peak to remain approximately constant. Finally, in model **B** zygosity shifts both the location of the two bifurcations and the level of the solutions, leading to noticeable changes in both expression level and the fraction of cells expressing the receptor. Similar bifurcation diagrams can be made for the *β* and *H* parameters, see supplementary.

Mathematically the source of bistability in all models is the nonlinear term. In the expression of the steady state solution of each model we can see the relation between copy number and the nonlinearity (Fig 4D). The gene copy number, *n*_*z*_, does not directly impact the nonlinear equation of model **A**, which explains the constant fraction of expressed cells exhibited by this model. The solutions will be multiplied by *n*_*z*_, which causes the shift in the expression level. In model **B**, in contrast, *n*_*z*_ affects both the nonlinearity (and thus the fraction of expressing cells) and the solution level (and thus the expression). Finally, in the nonlinear equation of model **C** the gene copy number only changes the basal production term. This changes the fraction of expressing cells but has a small impact on the high receptor solution, because *β*_*R*_ is large compared with *n*_*z*_.

**Fig 4.**
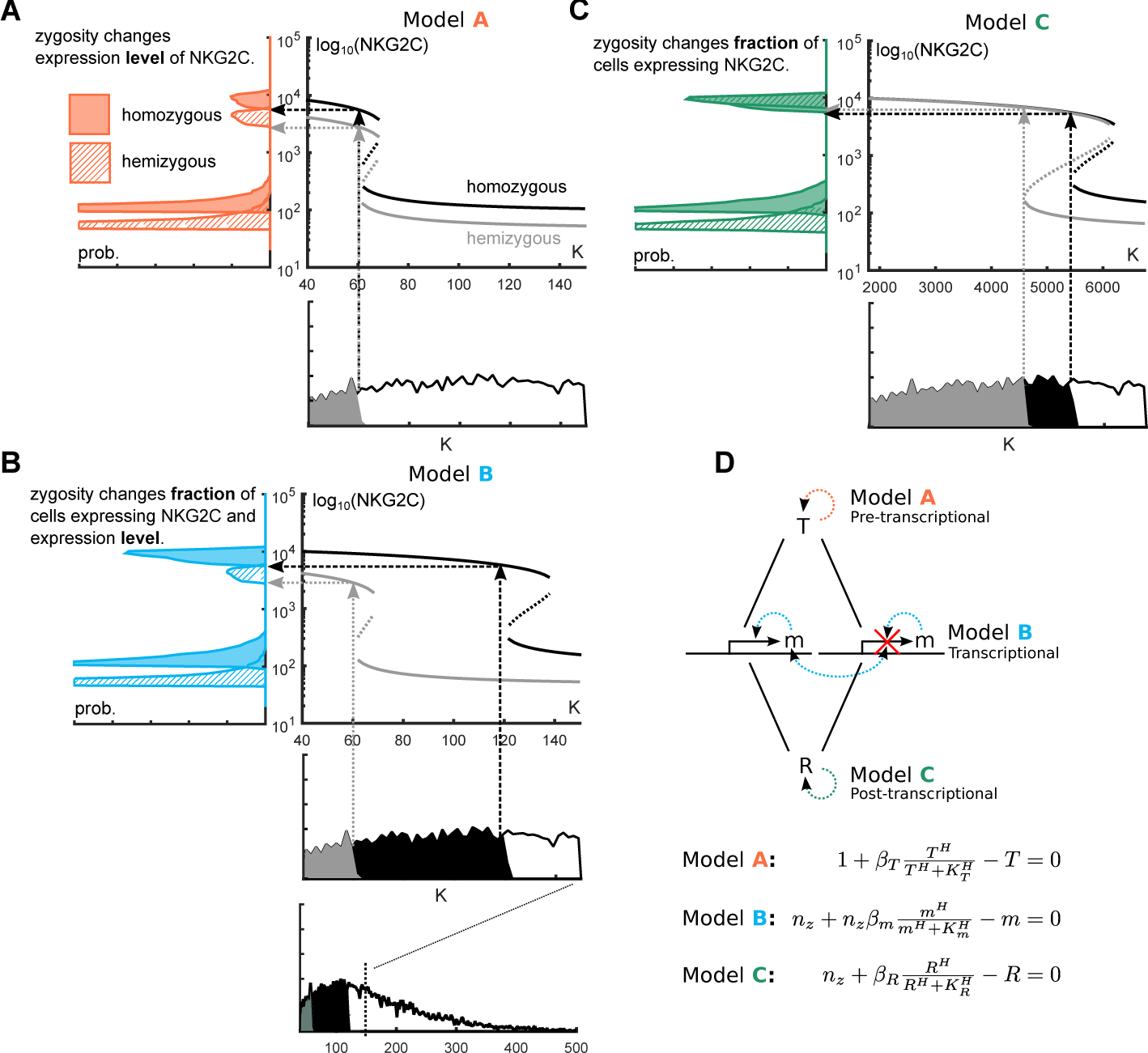
Quantitative model differences are explained by how zygosity influences bifurcations. NKG2C distributions as produced by noise in the *K* parameter. Each *K* parameter is translated through the bifurcation diagram into a receptor level. **A**: Model A (pre-transcriptional positive feedback): Zygosity does not shift position of the bifurcations in *K*. The solution values are however shifted by a factor two. **B**: Model B (transcriptional positive feedback): Zygosity changes both position of bifurcations and solution level. This translates into a change in both level and fraction of NK cells expressing NKG2C. **C**: Model C (post-transcriptional positive feedback): The bifurcation is shifted by a zygosity change while the high value solution falls on top of each other, i.e. the fraction is changed but not the receptor level. The *K* parameter is drawn from a gamma distribution with *µ*(*K*) = 160 and shape *θ*(*K*) = 60. **D**: Sketch of the regulatory network in each model. The nonlinear steady state equation is affected differently by *nz* in each model, leading to the different bifurcation diagrams.

In conclusion, model **B** is the only model that predicts both changes in expression level and fraction with zygosity, as observed among HCMV positive donors in Fig 1. The bifurcation diagrams further show that the location of the positive feedback relative to transcription gives three qualitatively different types of behaviour. Distributional changes due to zygosity can therefore help locate positive feedbacks in bimodal distributions.

### Discussion

The role of copy number variation in disease susceptibility and pathogenesis is becoming increasingly more recognized [21, 22]. However, the relation between gene copy number and phenotype expression is only partially understood. Inspired by an observed phenotypic difference between homo- and hemizygous HCMV positive donors in natural killer cells [13], we have shown that in the special case of a bimodal phenotype caused by a bistability, the expression differences due to zygosity can provide valuable information.

### Model validity and fitting

Within each donor group (zygosity and seropositivity) there were large variations in expression, ranging from no expansion to strong bimodal expression of the NKG2C+ NK cell subset. To test if the use of a median donor was reasonable under these conditions we did a cluster analysis (see Supporting Information). The analysis showed that seropositive donors can be divided into two subgroups: no expansion and bimodal distribution. The cluster analysis further showed that for donors with bimodal expression, the position of the second peak is a strong biomarker of zygosity. We have not shown any fits of single donors, but each model fits a large range of distributions reasonably well (see Supporting Information).

If we assume that not all cells in homozygous donors have two functioning genes, the model can be modified by introducing a subpopulation which effectively has one gene. This would create a superposition of a homo- and hemizygous distributions, which could help fitting (but also introduces an extra free parameter). Notice this does not change the results from the bifurcation diagrams, but zygosity effects will decrease as the fraction of cells with one functioning gene increases.

### Biological interpretation of positive feedbacks

Our modeling study suggests that, in order to reproduce the bimodality observed experimentally, the feedback has to include a transcriptional component. All the models considered here can be interpreted as single-cell models, but the processes are not necessarily restricted to individual cells. For instance, a bistable *T* variable could be an upstream transcription factor, but also a bistable external input. The *T*-variable can therefore be any or all processes upstream of transcription.

Model *C* is more restricted, as a post-transcriptional positive feedback can only be between transcription and the mature receptor in the cell membrane. Even though *R* is set equal to the membrane protein in this study, the positive feedback can be located downstream of transcription but upstream of the mature receptor. The effect would be the same if the positive feedback is followed by linear processes. A candidate for this type of behaviour could be the CD94 protein, which dimerizes with NKG2C to create the mature receptor [23]. Any bistability in CD94 would be reflected in the phenotype expression.

In model **B**, in turn, the positive feedback could in principle extend outside the cell as long as its impact reaches the production of mRNA in the end. It has been suggested that NKG2C and the inhibitory receptor NKG2A, which both dimerize with CD94, are mutually exclusive in CD8+ T-cells [24]. Another study has shown that NKG2C and NKG2A were reciprocally expressed in CD56dim NK cells, but co-expressed in CD56bright NK cells [25]. Receptors which mutually inhibit each others’ transcription is an example of a positive feedback at the transcriptional level.

Positive feedback can also arise at the population level. Previous studies [13] have shown that there is a correlation between proliferation rate and NKG2C level. This could lead, assuming that daughter cells inherit their mother’s receptor levels, to a positive feedback at a population level, since NKG2C-expressing NK cells will grow faster than other NK subsets. NK proliferation has previously been modeled [26], but not in a homeostatic state and not in relation to the NKG2C receptor. In the models presented in this paper proliferation is not taken into account. It is therefore not clear if NKG2C-dependent proliferation can be included in one of the models presented here, and proliferation-dependent bimodality could justify further investigation.

### Zygosity and gene copy number effects

Zygosity has previously been identified as a source of NK receptor alterations, but through its effect on the receptor ligand than on the receptor itself. Specifically, it has been shown that the zygosity of HLA-Cw7 (coding for NK receptor ligand) altered the NK CD158+ subset [27]. These alterations are therefore rather a response to changes in the target cells than a reflection of zygosity directly impacting phenotype expression. Another study has shown that increase in gene copy number of a NK receptor can have a positive impact on the expression of other receptors. In particular, the two genes KIR3DL1 and KIR3DS1, coding for killer cell immunoglobulin receptors (KIR), are important in the containment of HIV-I [28]. Similar to NKG2C, the KIR3DS1 receptor is only expressed by a subset of the NK population. Interestingly, the study by Pelak et al. [28] showed that the fraction of KIR3DS1+ cells and RNA transcription level increased with increasing copies of KIR3DL1. This resembles the observations from model **B**, except the changes are in response to copy number variation of a gene coding for another receptor.

In this study we have only considered one or two gene copies, but some NK receptors belonging to the KIR-family are observed to have three gene copies [29]. The mathematical analysis and the qualitative differences between models A, B and C will still be true for *n*_*z*_*>* 2. The aim of our models was to understand the differences between donors of different zygosity, but also to provide a rough tool for locating positive feedbacks of bistable phenotypes. Given a bistable phenotype, mathematical modeling can use donor groups of different zygosity to restrict the location of the positive feedback, as we have exemplified with the expression of NKG2C in HCMV positive donors.

## Methods

### Fitting and Model Comparison

Fit quality is quantified by Pearson’s *χ*^2^-value, which together with the number of degrees of freedom (ndf) provides a measure of the quality of the fit. We also use the Akaike information criterion (AIC) as a measure of relative model quality. The AIC value is given as: AIC = 2*k* − 2 ln(*L*), where *k* is the number of estimated parameters and *L* is the maximum value of the likelihood function. Assuming that residuals are distributed according to identical independent normal distributions, the Akaike Criterion can be rewritten to AIC = 2*k* − *n* ln(RSS*/n*), where RSS is the residual sum of square 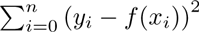. For small sample sizes the AIC is biased, but can be corrected by the addition of an extra term: 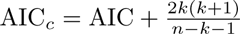 We calculate a combined Akaike score for each model using the results for both types of zygosity. The relative strength of evidence for each model is proportional to exp 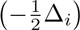), where ∆_*i*_ = AIC_*i*_ − AIC_min_. This can be summarized by a set of weights:

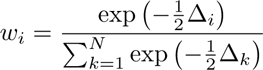

where 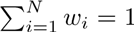. These weights determine the model that best represent the data relatively to the others.

## Acknowledgements

This work was supported by the Spanish Ministry of Economy and Competitiveness (MINECO) and FEDER (projects FIS2015-66503-C3-1-P and SAF2013-49063-C2-1-R), the Center for Models of Life, Niels Bohr Institute, Copenhagen University, and the Red Espan˜ola de Esclerosis Mu´ltiple (REEM, project RD12/0032/0005 ISCIII-MINECO). J.G.O. also acknowledges support from the ICREA Academia programme, the Generalitat de Catalunya (project 2014SGR0947), and from the “María de Maeztu” Programme for Units of Excellence in R&D (MINECO, project MDM-2014-0370). M.L.-B. and A.M. are also supported by the EU FP7-MINECO Infect-ERA program (project PCIN-2015-191-C02-01).

